## Supplementary figures and images for "SARS-CoV-2 disease severity and transmission efficiency is increased for airborne but not fomite exposure in Syrian hamsters"

### supplemental figure

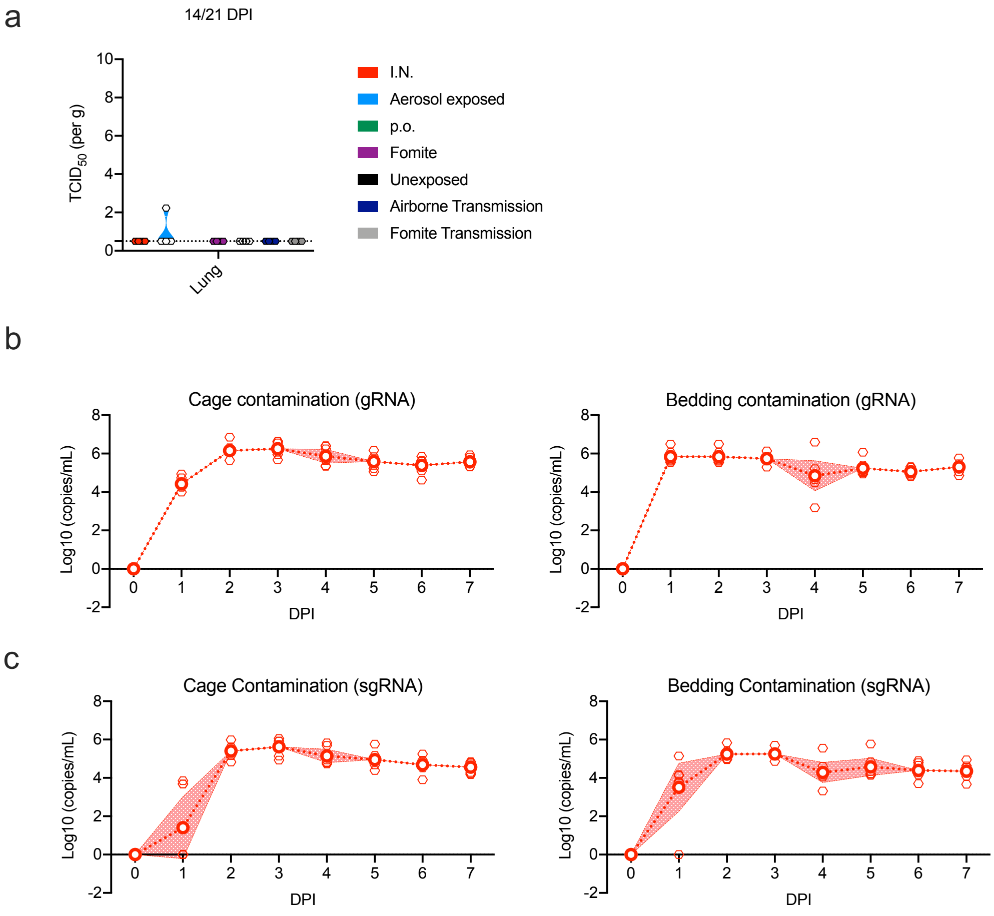
